# Hollow ring-like flexible electrode architecture enabling subcellular multi-directional neural interfacing

**DOI:** 10.1101/2022.12.24.521855

**Authors:** Venkata Suresh Vajrala, Kamil Elkhoury, Sophie Pautot, Christian Bergaud, Ali Maziz

**Author notes:** Corresponding authors. MEMS Group, LAAS-CNRS, 31031 Toulouse, France.

## Abstract

Implantable neural microelectrodes for recording and stimulating neural activity are critical for research in neuroscience and clinical neuroprosthetic applications. A current need exists for developing new technological solutions for obtaining highly selective and stealthy electrodes that provide reliable neural integration and maintain neuronal viability. This paper reports a novel Hollow Ring-like type electrode to sense and/or stimulate neural activity from three-dimensional neural networks. Due to its unique design, the ring electrode architecture enables easy and reliable access of the electrode to three-dimensional neural networks with reduced pressure on the biological tissue, while providing improved electrical interface with cells. The Hollow ring electrodes, particularly when coated with the conducting polymer PEDOT:PSS, show improved electrical properties with extremely low impedance and high charge injection capabilities, when compared to traditional planar disk-type electrodes. The ring design also serves as an optimal architecture for cell gowth to create an optimal subcellular electrical– neural interface. In addition, we demonstrated that the quality of recorded neural signals by the ring electrode was higher than recordings from a traditional disk-type electrode in terms of signal-to-noise ratio (SNR) and burst detection from 3D neuronal networks *in vitro*. Overall, our results suggest the great potential of the hollow ring design for developing next-generation microelectrodes for applications in neural interfaces used in physiological studies and neuromodulation applications.

## 1. INTRODUCTION

High-fidelity electrical interfaces for recording and stimulating neural activity are critically important for the purpose of understanding real-time dynamics of brain networks and managing neurological conditions. Current technological approaches and therapies, involve the use of implantable microelectrode arrays (MEAs) chronically interfaced with the neural tissue, possessing the ability to record from and stimulate small populations of neuronal cells. Such uni/bi-directional communication of neural interfaces combined with signal digitization, data processing and classification, has facilitated major advances on the treatment of various neurological conditions, such as deep-brain stimulation (DBS) for Parkinson’s disease^1-2^ and cochlear implants^3-4^, epilepsy^5^, depression^6^, chronic pain^7^ and other neurological disorders.

With the ever-expanding application of neural microelectrodes, one crucial challenge is to develop flexible microelectrodes that can closely record and/or electrically stimulate neural activity from small groups of neurons with high fidelity and reliability, without causing damage or altering other functionalities.^8^ Microelectrodes for neural recording transduce the bioelectric potentials carried from ionic currents into electrical signals through capacitive coupling. These electrodes should exhibit low impedance electrode/tissue interface to maintain signal-to-noise (SNR) quality for neural recordings on long term periods. For electrical stimulation, electrodes should be capable of injecting a sufficient and safe amount of current or voltage-controlled charge without causing any deleterious effects on either the electrodes or the surrounding tissue.^9^

Microfabrication technologies offers many advantages for the batch manufacture of reliable, microscale electrode array. Today, the most widely used and commercially available intracortical electrode arrays include thin-film planar (disk-type) electrodes based on silicon^10^, and bulk micromachined arrays.^11-14^ Such arrays have been used for localized recording and stimulation of neural tissue. Of these types of electrodes, planar electrodes, with the electrode diameter being in the range about 30-100 micrometres or larger, have been most broadly used to interface implantable devices with neural cells, thanks to their facile two-dimensional (2D) planar processing and practical use for *in vitro/in vivo* applications.^14-16^ However, conventional planar electrode configurations are not optimized for high-fidelity and long-lasting electrode-neuron interface. Small-size planar electrodes show low site-stimulation selectivity and poor resolution of cells, which can be attributed due to increased electrolyte resistance and shorter charging time of double-layer capacitance on electrode.^17^ This is light of the fact that neural cells are highly packed in three-dimensional assemblies of cell bodies, and a 2D planar electrode do not form a more realistic cytoarchitecture to access those soft and nonplanar cellular networks. Moreover, the reduction of electrode size at the single cell level, leads to higher interfacial impedance, thus the recorded SNR and stimulation efficiency will be reduced.

In this regard, the selection of electrode geometry and electrode coating material were found to have significant impact on enhancing the electrodes capabilities i.e., electrochemical impedance and charge transfer capacity. Several novel curved and folded electrode configurations were reported in the literature, such as nanowires^18^, star-shaped^19^, honeycomb-shaped^20^, spiral shaped^21-22^, spine-shaped^23^, shell-shaped^24^, cylindrical shaped electrode arrays^25^, to improve the neural surface interfacing capabilities. These unconventional electrode geometries offer higher perimeter-to-surface ratio, compared to the regular disk-type electrodes and exhibit minimal access resistance and increased ion flux to the electrode interface.^26^ Hence, it is an effective way of reducing the interfacial impedance and improving charge injection limits. However, most of these electrode configurations were built exclusively for only *in vitro* applications where the recording area would be mounted from the bottom of the cell cultures, where it is attached to the plate. In addition, a few of these propositions mainly focused on numerical analyses without providing systematic electrochemical and electrophysiological investigations. In parallel to the electrode geometry, new strategies utilizing rough organic conductive nanomaterials such a conductive polymers (e.g. PEDOT:PSS)^27-28^, carbon nanotubes (CNTs)^29^, carbon nanofibers (CNFs)^30-31^, graphene^32^, reduced graphene oxide (r-GO)^33^ and their nanocomposites^34-36^ have allowed to improve electrochemical surface area, while still providing the desired geometric area.

Whereas on other hand, in the context of implantable neural electrodes devices, the problem of large mechanical mismatch is still existing at the neural interfaces. Immune reaction occurs through a serious of mechanical compression events between rigid planar electrodes e.g., glass, silicon, gold or platinum, constituting the electrodes and the softness of tissue, triggering inflammatory reactions and glial scar generation, which directly impact on the survival of both electrode and neurons.^37^ These device failures lead to higher interfacial impedance, which impact the recorded SNR and the stimulation efficiency. Strategies to minimize these adverse tissue reactions revolve around the use of soft biomaterials as substrates such as parylene^38-39^, polyimide^40^, silk^38^, polydimethylsiloxane (PDMS)^41^ or hydrogels^42^, to fabricate flexible neural interfaces, to achieve better compliance with the soft neural tissue with minimal inflammatory reactions.^43^

Overall, a need exists to develop flexible devices with an improved electrode geometry that would allow an intimate contact with neurons and delivers reliable neural interfacing capabilities. Although incremental progress in this area has been substantial^44-46^, there is still a room for improving definitive technological solutions for obtaining highly selective and stealthy electrodes that provide reliable neural integration and maintain neuronal viability. A global rethinking of novel materials, geometries, and fabrication strategies is therefore needed for developing the next generation of neural interfaces that will overcome those issues. In this work, we show that it is possible to tackle several of those above-mentioned challenges by the development of a novel electrode architecture that may lead to an improved tissue integration, while still providing optimal electrical characteristics to control intrinsic biological processes. We describe the design and characterization of Hollow-Ring-shaped electrodes (Figure 1), coated with nanostructured PEDOT-PSS, fabricated on a non-invasive flexible parylene backbone, for improved signal recording and electrical stimulation. The Hollow-Ring-shaped design permits the dynamic adaptation of the electrode to the chronic changes in the 3D neural network. The ring-shaped electrode architecture with microscale geometrical surface area (smaller electrode footprint and higher perimeter-to-surface ratio) were designed specifically in such a way that it offers more freedom to the internal dynamics of neural networks, at the same time offering multidirectional sensing of neuron activities within the 3D neuronal networks.

**Figure 1:**
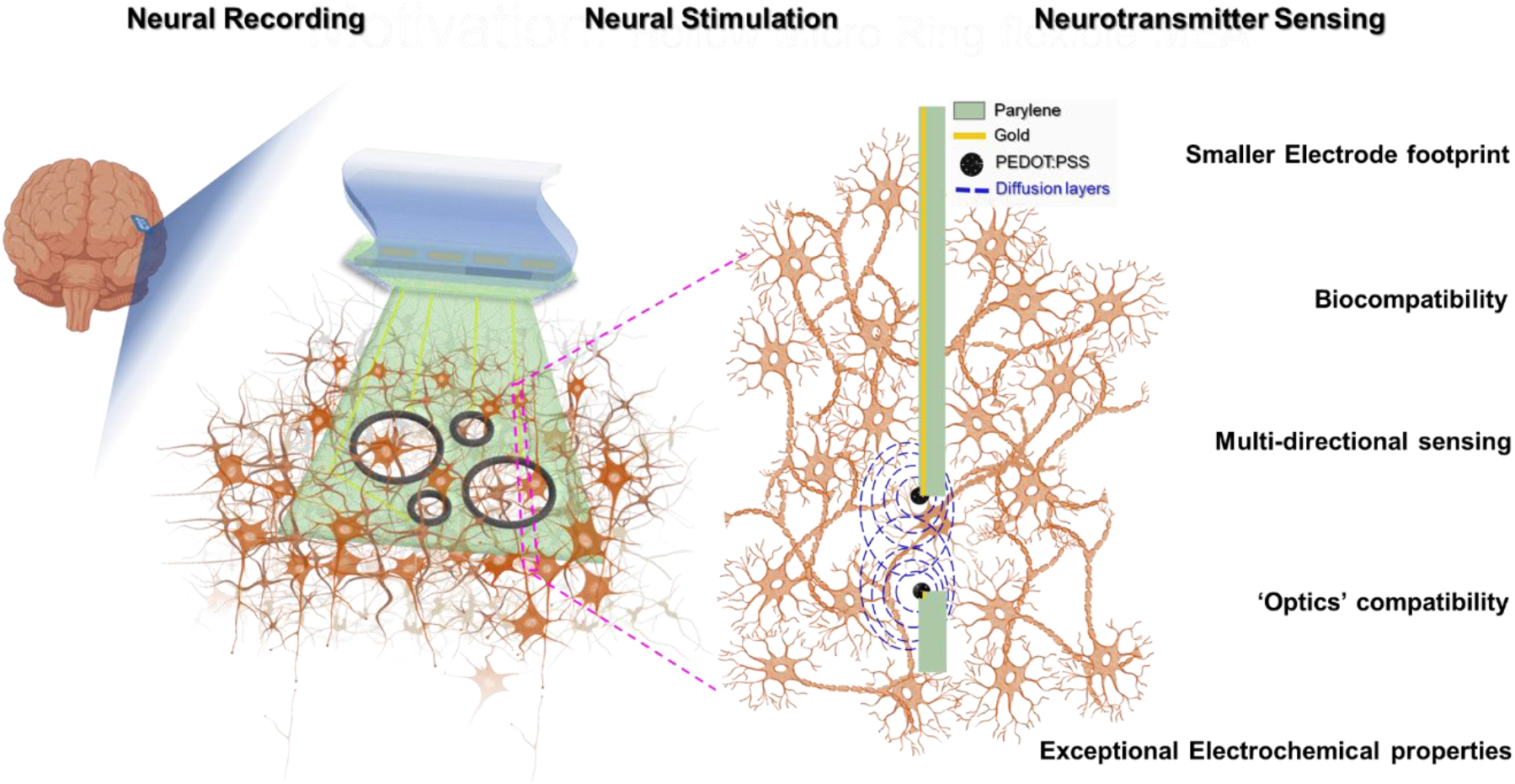
Schematic illustration of Hollow-Ring-type flexible microelectrode having different hole sizes, inserted within neural network, and their potential features and applications.

## 2. MATERIALS AND METHODS

### 2.1 HR-microelectrodes fabrication process

The design and fabrication process of Hollow Ring (HR) microelectrode array was illustrated in Figure 1, Figure 2a & b, respectively. In brief, a 10 μm-thick film of parylene C was chemically vapour deposited (Comelec C-30-S at 700 °C) onto a clean 4-inch” SiP wafer. Subsequently, gold disk type microelectrodes (Ti 50 nm/ Au 200 nm) were patterned on the parylene surface by physical vapour deposition followed by a lift-off process with AZ-nLof photoresist (AZ-nLOF 2035, MicroChemicals). Next, a passivation layer of 3 μm thick parylene was deposited and annealed at 110°C for 16 hours under N_2_ flow. The active electrode areas were then realised, by pattering with 5 μm-thick AZ4562 photoresist (MicroChemicals) film, followed by the removal of thin parylene layer using O_2_ plasma reactive ion etching ICP-RIE (Trikon Omega 201). After that, the connecting pads and outlines of implant were realised, by patterning with 50 μm-thick BPN photoresist (Intervia BPN-65A, Dupont) and etching anistropically using deep reactive ion etching (ICP-DRIE). Later, the implants were released out of the silicon substrate in DI water, stripped off from the remaining resist using TechniStrip NF52. The implants were bonded to flexible ribbon cable (blue color) with golden traces finally, as shown in the Figure 2a. At the end, the surfaces were cleaned by repetitive cycles of ethanol (99,5%; Fisher Scientific) and DI water, for 10 minutes each cycle and stored in a dry place.

**Figure 2:**
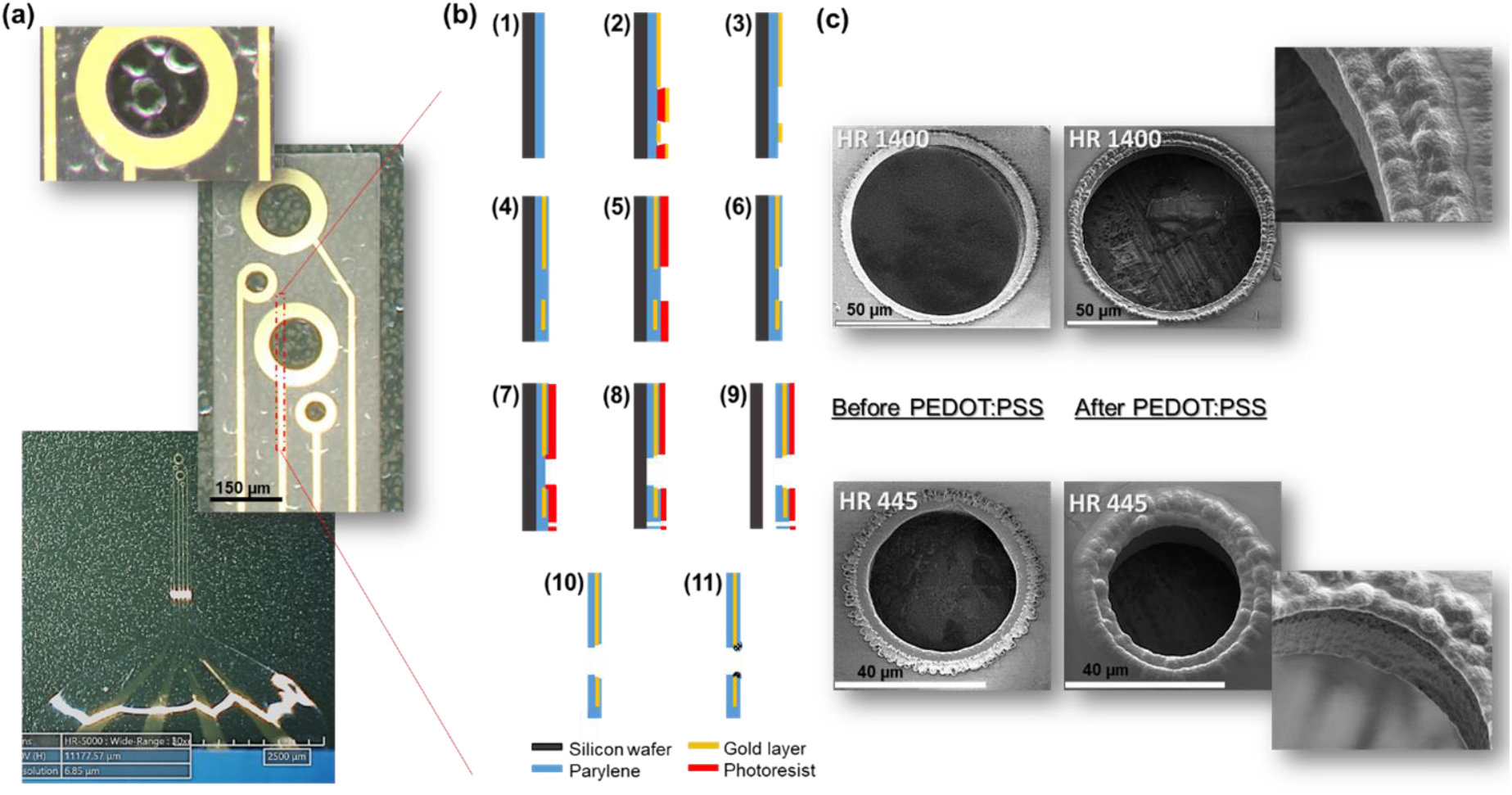
(a) Hirox microscope images of Hollow Ring microelectrode array. (b) Schematic overview of steps involved in the microfabrication procedure- (1) 10 μm thick Parylene C deposition on SiP wafer, (2) micropatterning of nLof photoresist and gold layer deposition (Ti/Au-50/200 nm), (3) nLof removal and cleaning, (4) deposition of second Parylene layer of having 3 μm thickness and anneal at 110°C for 16h, (5) Photopatterning with AZ4562 photoresist, (6) RIE with ICP-RIE oxygen plasma, (7) Photopatterning with BPN photoresist, (8) RIE with ICP-DRIE oxygen plasma, (9) Stripping of implants from SiP wafer, (10) Development with NF52 and cleaning with DI water, (11) Electrodeposition of PEDOT:PSS on the ring type gold electrode. (c) SEM micrographs of Hollow ring microelectrodes, before and after PEDOT:PSS deposition.

### 2.2. PEDOT: PSS deposition and electrochemical characterization

The PEDOT: PSS deposition solution was prepared by mixing both 0.1% w/v NaPSS (Sigma Aldrich MW∼ 1000000) and 10 mM EDOT (Sigma Aldrich 97%) in DI water and stirring it for 2 hours at room temperature. Later, the deposition solution was purged with N_2_ and stored at 4°C until further use.

The surfaces of all gold electrodes (HR and Disk) were activated in 0.5 M H_2_SO_4_ by applying multiple electrochemical oxidation-reduction cycles, with the applied potential in between - 0.4V and 1.6V (vs Ag/AgCl electrode as reference and platinum wire as counter electrode), at a scan rate of 200 mV/s using Biologic potentiostat (BioLogic VMP3) (Figure S1). Upon the surface activation, the electrodes were rinsed in DI water and dried properly. Later, the deposition of PEDOT: PSS on all electrodes was conducted either using cyclic voltammetry (CV) or chronopotentiometry (CP) technique. For CV based deposition, every electrode was subjected to one CV scan cycle between -0.7 V and 0.9 V, at a scan rate of 10 mV/s vs Ag/AgCl electrode as reference. In CP technique, the PEDOT: PSS was deposited galvanostatically by applying a small current of 12.5 nA, with a constant charge density of 4 nC/μm^2^. After the PEDOT: PSS deposition, the electrodes were rinsed in DI water and immersed in PBS buffer for at least 30 minutes, before performing characterization.

To demonstrate the improved electrical functioning of electrodes, standard electrochemical characterizations, such as charge storage capacity (CSC), electrochemical impedance spectroscopy (EIS) and charge injection limit (CIL) were conducted. EIS and CSC measurements were performed in three electrode configurations, using a Bio-Logic VMP3 potentiostat, where Ag/AgCl as reference and platinum mesh as counter electrodes. Cathodic CSC values of all electrodes were estimated by launching a series of CV based sweeps on a low current potentiostat channel (BioLogic VMP3), between 0.7 V and -1.0V in PBS buffer at room temperature (Figure 3b). Every electrode was swept for 4 CV cycles and the CSCc was calculated as the time integral of the cathodic current recorded over a potential range of 0.6 V to -1.0 V in the last cycle. Impedance (EIS) measurements were conducted by applying 10 mV AC signal amplitude in the frequency range from 0.5 HZ to 5 MHz on all electrodes (Figure 3 a and Figure S3a). The water reduction potential was determined by performing a CV between -1.5 V to 0.7 V at 200 mV/s vs Ag/AgCl reference (Fig S3b & c). Biphasic, cathode first, voltage transient measurements were performed in PBS (pH 7.4) at a frequency of 10 Hz and with different pulse width durations ranging from 0.5 to 5 ms (Figure 3c & d and Figure 4). Different input current values were injected progressively, starting from 0.5 μA and reached up to 360 μA, depending upon the experimental condition, using a Bio-Logic VSP3 potentiostat and the resulting voltage excursions were measured. The negative polarization potential (V_p_) was calculated by subtracting the initial access voltage (V_a_) due to solution resistance from the total voltage (V_max_). The CILs were calculated by multiplying the current amplitude and pulse duration at which the polarization potential reaches the water reduction limit (−1.15 V), divided by the geometric surface area of the electrode. All electrochemical measurements were performed in a Faraday cage.

**Figure 3:**
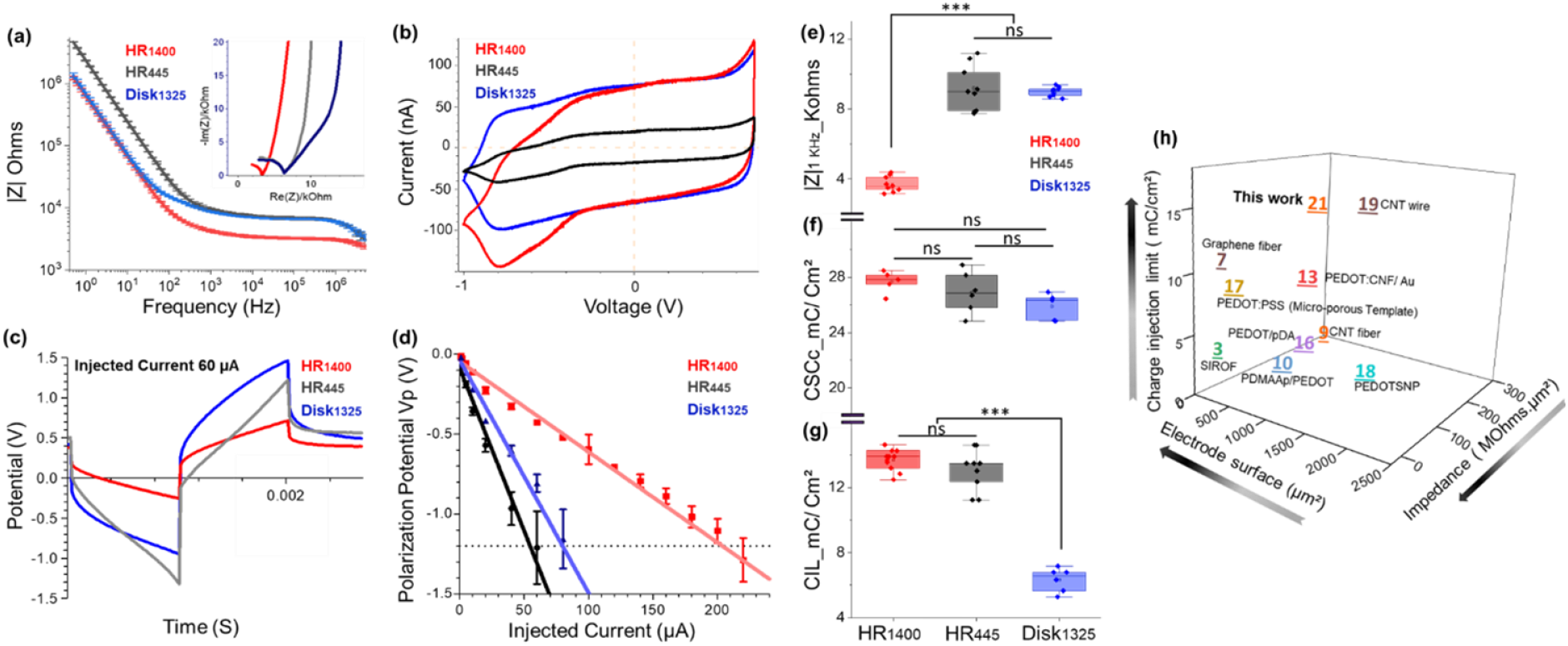
Electrochemical characterization of HR and Disk type electrodes coated with PEDOT:PSS (a) Bode plot representing the |Z| vs frequency over a frequency range of 0.5 Hz to 5 MHz in PBS at 0V vs Ag/AgCl ref electrode (Inset: Nyquist plot measurements obtained by EIS). (b) CSCc measurements by CV in PBS at 200 mV/s vs Ag/AgCl ref electrode. (c) Biphasic charge-balanced voltage responses after injecting the electrodes with the current of 60 μA. (d) Polarization potentials (V_p_) measured under different current pulse amplitudes. The box plots representing the evolution of (e) |Z|1kHz (f) CSCc (g) CIL values and the statistical differences between electrode responses were assessed by ANOVA followed by a simple t-test, where *** and ns represent P<0.001 and no significant difference, respectively (n=4). (h) 3D graph representing the comparison of Impedance, CIL performances and geometric surface area of HR microelectrodes coated with PEDOT:PSS with other neural electrode arrays reported in the literature (Table 1).

**Figure 4:**
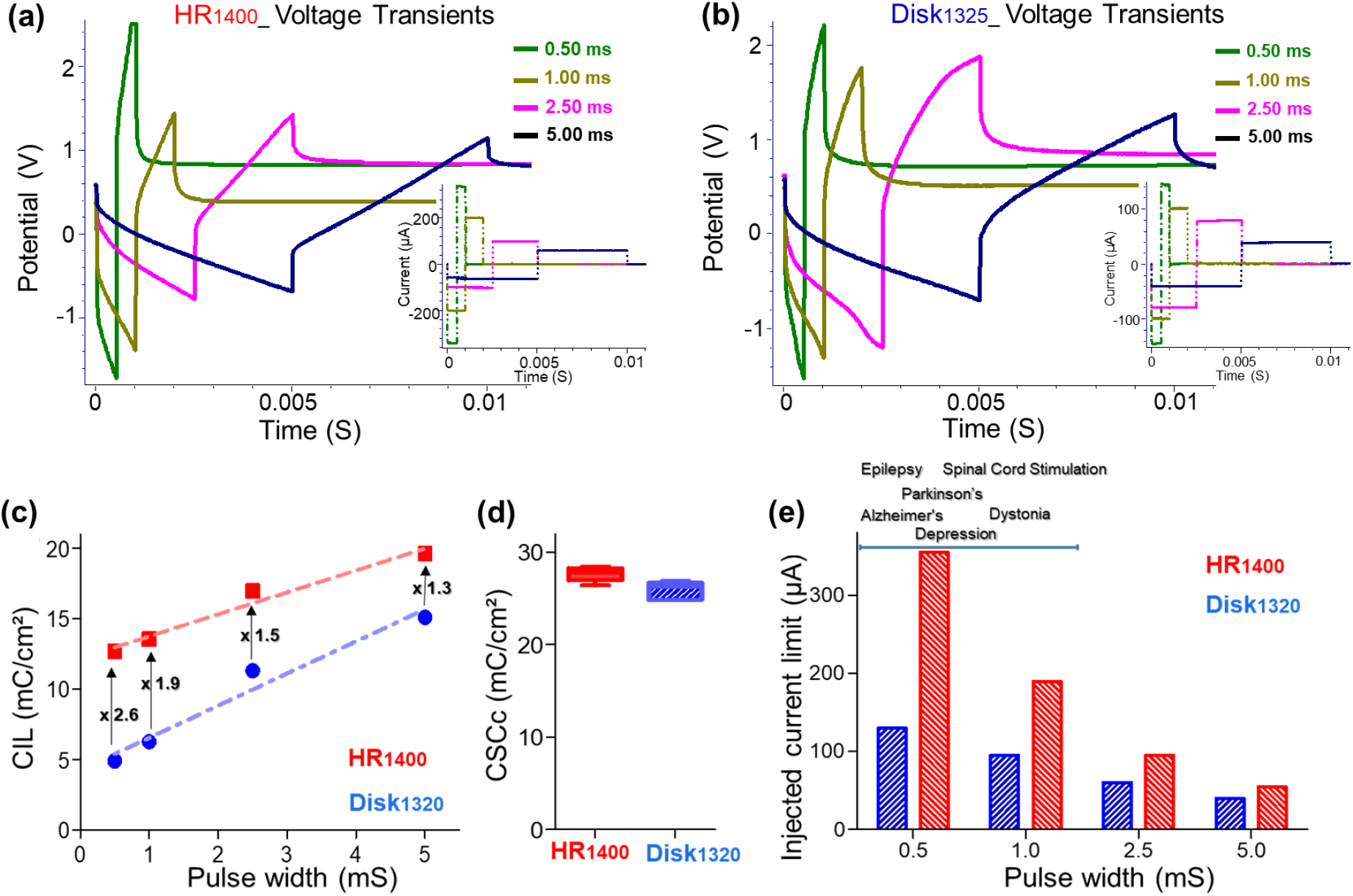
Biphasic stimulation performances of HR and Disk electrodes at different pulse widths ranging from 0.5 ms to 5 ms. Biphasic charge-balanced voltage responses of (a) Hollow ring, (b) Disk type electrodes and their respective injected current pulses were shown in the insets. (c) Polarization potentials (V_p_) measured under different pulse widths. The arrows at each condition describe the amount of CIL increment observed. (d) Comparison of average CSCc values and (e) maximum injected currents, before reaching the water reduction limit, of both HR and Disk type electrodes.

### 2.3. SH-SY5Y cell culture and viability tests

#### SH-SY5Y cell culture

The human neuroblast cell line (SH-SY5Y) was obtained from the American type collection. Dulbecco’s modified Eagle’s medium containing glutamax, pyruvate (Gibco, Invitrogen) and 10% fetal bovine serum was used for the SH-SY5Y. Cells were seeded at a density of 15000 cells/cm^2^ in 24-well plates w/wo implants and incubated for 1, 3, and 7 days at 5% CO2 and 37 °C. Prior to seeding implants were sterilized in 70% ethanol for 30 mins.

**Table 1:**
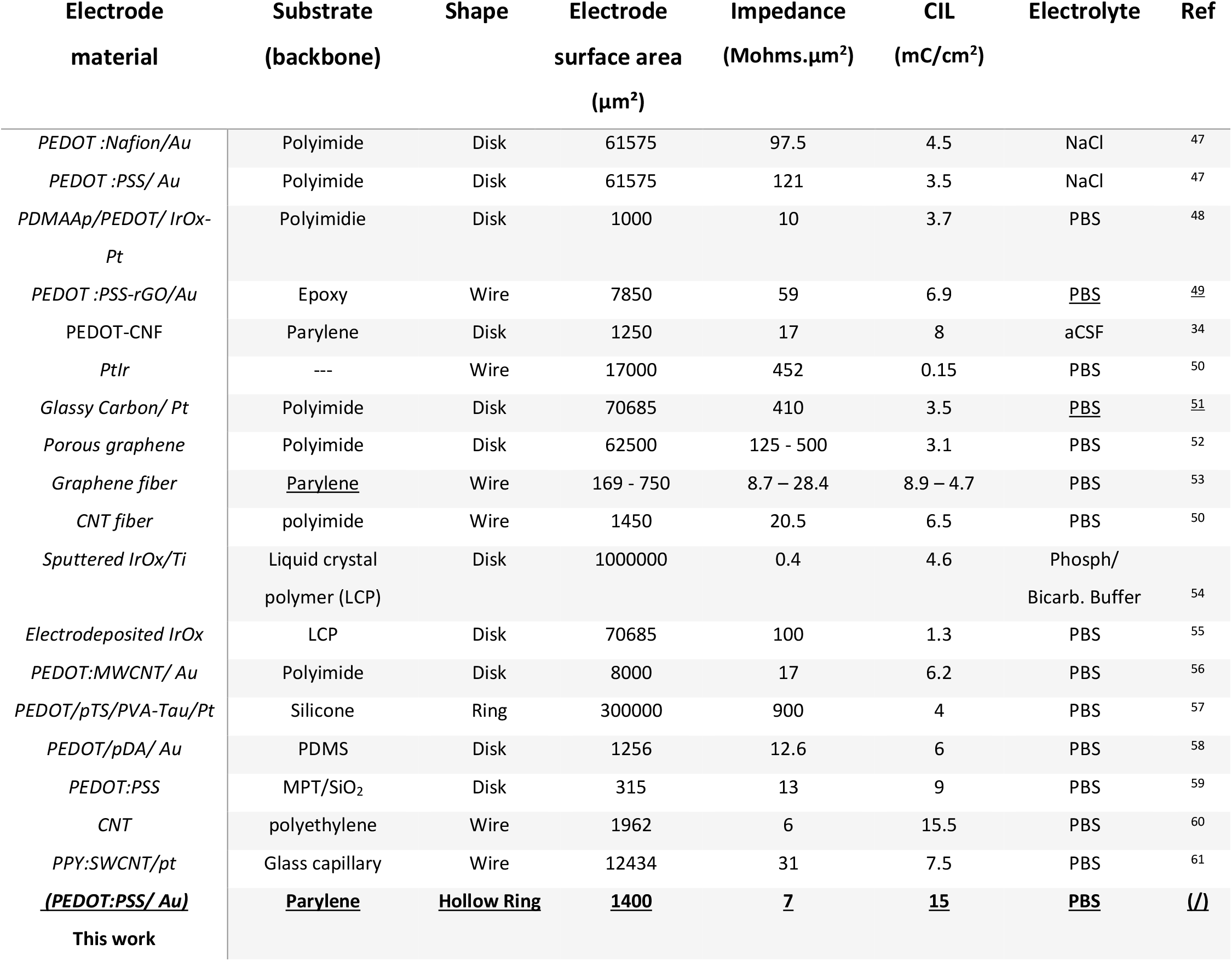
Comparison of the electrochemical performances of our HR microelectrodes coated with PEDOT:PSS with few of the well-known materials to fabricate neural interfacing microelectrodes.

#### Live/Dead assay

A mixed solution containing 2 μL/mL of Ethd-1 and 0.5 μL/mL of Calcein AM (Sigma-Aldrich) was prepared. The solution was added to the cells after media aspiration, and the cells were incubated for 30 mins at 5% CO2 and 37 °C. After incubation, cells were washed two times with PBS, and images were taken using a fluorescent microscope.

#### MTT test

At day 1, 3, and 7, cells were incubated for 2 h at 37 °C and 5% CO_2_ with MTT (Sigma-Aldrich) dissolved in DMEM media without phenol red. Culture media was then replaced with DMSO for 30 mins at 37 °C and 5% CO_2_. Finally, absorbance was measured at 570 nm.

### 2.4. Fabrication of cell culture chamber and packaging of HR microelectrode array

Cell culture chamber and HR microelectrode array holder (lid and arm) were designed using AutoCAD and printed using a 3D printer 29J+ HR (DWS, Italy). The cell culture chamber (*d* 2.6 cm and *h* 1 cm) was mounted upon a glass coverslip using PDMS as adhesive. The holder lid was printed using acrylate resin DS3000 (DWS, Italy) and 405 nm laser at a writing speed of 5.8 m/s. It was designed to hold the HR MEA and align vertically to the glass MEA, as shown in the Figure 6a.

### 2.5. 3D Neuronal culture and silica bead preparation

Borosilicate glass spheres (45 μm *d*, MO-SCI Speciality Products) and all components of the MEA setups were sterilized overnight in ethanol and dried under vacuum. Later, the beads were incubated in borate buffer for 1 h, followed by an overnight incubation step with poly-l-lysine (PLL) solution. Meanwhile, hippocampus was removed from rat embryonic day 18 embryos (E18), treated with trypsin and then triturate into a cell suspension, with a fire-polished glass Pasteur pipette.

The cells were seeded at 75000 cell/ cm^2^ on PLL coated glass beads and later, they were dropped onto bare coverslips upon which the monodispersed beads assembled spontaneously arranged into 2D ordered arrays. Once the neurons reached maturity on their respective beads, we carefully moved the beads carrying the cells to the cell culture chamber mounted with both HR and Disk type microelectrode arrays with a coarse pipette. The beads settled under gravitational force and spontaneously assembled into 2D hexagonal arrays at the bottom of the chamber. Once the first layer was fully packed, more batches of beads were added successively resulted in a construction of 3D packed assembly over the microelectrodes. The beads were arranged together to form stable hexagonal packing structure, eliminating the need for exogenous cross-linking agents to hold the arrays together during solution exchange. In addition, the void spaces between the beads permitted the necessary medium exchange to maintain healthy growing conditions, without disrupting their membrane integrity. Cells were cultured in neurobasal medium (2 mM Glutamax, 2% FBS and 2% B-27) for at 37°C. The neuron maturation starts from the 3^rd^ week and the samples were kept in the incubator, following 3 months without further shaking, but with regular medium exchange.

### 2.6. In vitro electrophysiological monitoring and pharmacological testing

A wireless W2100-HS4 electrophysiology system (Multichannel Systems, Reutlingen, Germany) was used to record electrophysiology activity from flexible Hollow-Ring and Disk type microelectrode devices for 30 min at an acquisition frequency of 20 kHz and bandpass filtered between 200 Hz – 3000 Hz. The cell culture chamber, attached with HR MEA system, was placed within a 5% CO_2_-regulated chamber on a heated stage at 37°C. Before recording spontaneous activities of neurons, the system was left alone for 10 mins to establish equilibration time. A software-supplied spike detector was used to detect spontaneous events that exceeded a threshold of ±4 μV (an extracellular action potential spike was defined by a lower limit threshold, set at 4.5× the standard deviation of baseline noise, for each electrode) (Figure 6d & f). We also monitored low frequency busts events using a band-pass filter 0.1 Hz – 30 Hz (Figure 6e & g). Data was exported as a hdf5 file and analysed using an in-house custom R package to calculate electrophysiological features such as number of spikes and bursts. Burst parameters, defined previously include maximum beginning ISI of 0.1 s, maximum end ISI of 0.2 s, minimum IBI of 0.5 s, minimum burst duration of 0.05 s, and minimum number of spikes per burst of 6. Features were calculated per channel for each device. The mean standard error of the mean (SEM) values per device were calculated for each feature.

### 2.9. Statistical Analysis

The statistical analyses were performed by Origin software (OriginPro 8.5, USA) and all the conditions were statistically evaluated by one-way analysis of variance (ANOVA) with t-test (p < 0.05), where ** = p < 0.01, *** = p< 0.001 and NS = no significant difference.

## 3. RESULTS AND DISCUSSION

### 3.1. Design and fabrication of flexible HR microelectrode array

The main goal of this research is to design and fabricate a novel neural electrode that can offer multisided operability, while offering superior electrode properties and smaller footprint of a functional electrode surface, i.e., exposed to electrolyte media. In this regard, we fabricated a novel Hollow-Ring type gold microelectrode array on a flexible parylene backbone, having a thickness of 13 μm. It offers a unique rounded design space for neurons to envelope and allows a multidirectional measurement capability of electrophysiological responses that is not accessible with conventional 2D disk planar electrode surfaces.

As described in Figure 2a & b, we followed standard photolithographic and reactive ion etching procedures to achieve customizable well-defined ring-like electrode structures, patterned on a flexible and biocompatible parylene backbone.^34-35, 38-39^ The photomask designs and etching parameters were optimised carefully during the microfabrication process, so that the gold electrode traces were revealed while avoiding the over etching of both parylene layers, which could enlarge the electrode surface area. All the four electrodes in HR microelectrode assembly were spanned in one direction and each probe consists of two different inner ring diameters (IRD) and surface areas (S.A), i.e., HR_445_ (40 μm IRD and S.A. 445 μm^2^) and HR_1400_ (100 μm IRD and S.A.1400 μm^2^). The intended designs of the inner ring electrode diameters (40 μm and 100 μm) and their respective surface areas (445 μm^2^ and 1400 μm^2^) of our HR microelectrodes were deliberately crafted, for comparison purposes, where they match with the physical dimensions of the traditional disk type electrodes, having 40 μm electrode diameter, with the surface area of 1325 μm^2^.

Before post-processing step, all the electrodes were repetitively cycled in 0.5 M H_2_SO_4_ using cyclic voltammetry to obtain clean gold electrode surfaces. Later, PEDOT:PSS layer was electrodeposited on all electrodes to improve the signal stability and quality.^27-28^ PEDOT: PSS is a widely used conductive polymer that offers synergistic ionic/ electronic conductivity and inherently flexible biocompatible surface to create a seamless bridge between electrode and tissue interface. PEDOT:PSS films were deposited on gold electrode surfaces in aqueous solutions of 10 mM EDOT monomer and 1% w/v NaPSS, using galvanostatic and cyclic voltammetry methods (Figure S2). SEM observations suggested that the electrodeposition parameters critically affected the overall structure of the electrode surfaces. In the case of HR microelectrodes, the cyclic voltammetry-based depositions produced uncontrolled ‘balloon-like’ nodular rough surface morphology, extended away from the gold electrode, with few cracks on them (Figure S2b). Nonetheless, ensuring crack-free and well-controlled depositions were important in order to improve the deposition stability and signal quality over long-term. On the other hand, galvanostatic deposition method offered superior homogeneity and quality of the PEDOT:PSS layers over other method (Figure S2c). The Figure 2c, shows the galvanostatic deposition of PEDOT:PSS on HR microelectrodes, under the optimised current density of 10 pA/μm^2^ and surface charge density of 4 nC/μm^2^, where highly ordered depositions were observed uniformly spreading all around the hollow ring structures.

### 3.2. Electrochemical characterization

The elevated electrochemical impedance and inferior charge injection limit (CIL) values often become hurdles for the better performance of neural microelectrodes. Hence, an effective and stable information transfer of the neural interface is important. Microelectrodes with low impedance values favours monitoring the electrophysiological signals with better resolution. Simultaneously, a very high and stable charge injection limits enhance the electrical stimulation capability of the electrode, where it is defined as the maximum charge density that can be injected into the tissue before reaching the water reduction potential.

#### 3.2.1. Electrochemical impedance Spectroscopy (EIS)

The EIS measurements were carried out by sweeping over frequencies ranging from 0.5 Hz to 5 MHz. Typically, the neuronal frequencies reside within different frequency bands, but the impedance at 1 kHz is an important indicator relating to the frequency of neuronal single-unit recordings and power consumption during electrical stimulation. As shown in the Figure 3a and Figure S3a., the impedance magnitude at 1 kHz (|Z|_1kHz_) against frequency decreased sharply from uncoated gold electrodes to PEDOT:PSS coated electrodes and it was clearly the common trend for all the three electrode configurations (HR_1400_, HR_445_ and Disk_1325_) involved in this study (Figure S 4b). After depositing PEDOT:PSS at the same charge deposition conditions (4 nC/μm^2^) among three electrodes, HR_1400_ showed significant reduction in the impedance at all frequencies where the value of |Z|_1kHz_ being 3.7± 0.42 kΩ. Despite having comparable surface areas, the impedance magnitude of Disk_1325_ was found to be 10 times higher than HR_1400_. Whereas HR_445_ electrode, with almost 60 % less surface area than conventional Disk_1325_, exhibited similar impedance patterns and matched to the characteristics of Disk_1325,_ especially at higher frequency range (Figure 3a & e).

It is a well-known fact that the overall reduction of the electrical impedance is a result of expanded electrode surface area coated with the PEDOT:PSS film, where it establishes low ion transfer resistance at the electrode/electrolyte interface. However, by looking at both Figure 3a and Figure S3a, in addition to the smooth interface of electrolyte and polymer, it appears that the shape and the amount of electrode surface that is available to the electrolyte might influence the impedance magnitude.

To understand this phenomenon better, the Nyquist plot characteristics were analyzed as shown in the Figure 3a inset. A semi-circle region at higher frequency is ascribed to the charge transfer resistance at the electrode concomitant with the formation of the electrical double layer. Followed by a line with slope at low frequency region related to the Warburg resistance, reflecting the frequency dependence of ion diffusion/transport in the electrolyte. Larger the Warburg region, in x-axis direction, greater the variations in ion diffusion path lengths, hence the increased resistance to ion transfer. We observed a significant reduction in the charge transfer resistances (reduced semi-circle regions) and short Warburg path lengths (almost straight lines) corresponding to improved diffusion rates in HR electrodes compared to the conventional Disk electrodes. In addition, increasing the diameter of hole from 40 μm (HR_445_) to 100 μm (HR_1400_) significantly reduced the resistance to the charge transfer, which translates into an increased the surface area availability for the electrode-electrolyte interface and enhanced rate of charge transfer with lower impedance at higher frequency range.

#### 3.2.2. Cyclic Voltammetry

Under the optimized charge deposition density conditions (4 nC/ μm^2^), the PEDOT:PSS was deposited galvanostatically on HR and Disk type electrodes. The cathodic current was integrated from their respective CVs (Figure 3b), to define the CSCc values and we observed that the PEDOT:PSS deposition process on the gold electrode surfaces improved the CSCc thanks to the increased electroactive surface area (Figure S4c). It was hypothesized that the HR electrodes, especially HR_1400_ with its 100 μm hole in the middle, would possess higher CSCc due to the increased availability of active surface area, however, this was not observed. Figure 3f, showed that all three electrodes, after PEDOT:PSS coating, exhibited similar (statistically insignificant) CSCc values (25-30 mC/cm^2^). It indicates that the redox activity of PEDOT:PSS, at slow linear sweep rates, is less affected by the electrode shape, but relies more on the effective surface area and robust charge deposition density conditions. It is already a well-known fact that the values of CSCc might not be accompanied by the corresponding improvements in the charge injection limit, especially at high-frequency pulsed stimulation conditions, only a part of the CSC is accessible due to the limited diffusion rate of the charge carriers in the solution phase. Therefore, the charge injection limit (CIL) is a more practical parameter than the CSC in comparing the stimulation performance as neural electrodes, where CIL is measured using high-frequency current pulses.^8^

#### 3.2.3. Charge injection limit

The voltage excursions and the respective CILs of all electrodes were evaluated using cathodic-first biphasic current pulses having 1000 μs pulse width and 50 ms inter-pulse duration in PBS buffer (Figure 3c & d). The water reduction potential for HR and Disk electrodes coated with PEDOT:PSS were found to be around -1.2 V to -1.4 V (Fig S3c). We considered -1.15 V as the limit to stay safe and prevent overestimation of the CIL values.

As shown in Figure 3c, despite having comparable electrode active surface areas, the voltage excursion and the waveform of HR_1400_ showed very less polarization compared to Disk_1325_, when applying same current pulse (60 μA). The linear regression curves of polarization potential (V_p_) at different current pulses showed that HR_1400_ can hold the current pulses of at least 2-fold higher than the Disk_1325_ electrode, before reaching the water reduction potential (Figure 3d). These observations indicate that HR electrode utilizes nearly 50% of CSC during stimulations whereas for the Disk electrode, only 25% of the CSC is accessible. It is likely due to the limited diffusion rate of the charge carriers in the solution phase.

In addition, the CIL that was calculated at V_p_ = -1.15 V, before the water reduction potential (Fig S3c), to be 15 ± 2 mC/ cm^2^ for the HR_1400_ electrode; a value ≈ three orders of magnitudes higher than bare gold ring electrode and ≈ two times larger than the Disk-type PEDOT-coated electrode. The CIL of the HR electrodes was significantly higher than all the best reported electrode designs, including but not limited to disk, wire, or fibers coated or non-coated with carbon nanotube fibers, conducting polymer coatings, metal nitride and oxides, as presented in Figure 3h and listed in Table 1.

Furthermore, to understand the influence of electrode shape, we investigated the effect of pulse width on CIL for safe electrochemical stimulation. The shorter pulse widths in general increase the selectivity for large, distant axons, while long pulse widths focus on the stimulation effect on small, nearby axons^62^. Figure 4a & b, shows the voltage transients and the measured CILs of PEDOT:PSS coated HR_1400_ and Disk_1325_ electrodes, under various pulse widths ranging from 0.5 to 5 ms. Lower values of CILs were obtained with shorter pulse widths, because the entire active region of the electrode was not utilised due to the non-uniform current distribution. With longer pulses, the reaction has the time to spread all over the electrode surface to produce higher charge injection limits. The CILs at every pulse width condition showed that our HR_1400_ electrode outruns the traditional disk electrode, despite having comparable electroactive surface areas and CSCc values (Figure 4c & d). Whereas, by increasing the pulse width from 0.5 ms to 5 ms, the net CIL improvement was observed to be higher in the disk electrodes (∼ 175 %) than HR electrodes (∼ 50%). We also observed that, at lower pulse width conditions, the HR electrode can withstand almost 2 to 3 times higher injected currents than the disk electrodes, before reaching the water reduction potential (Figure 3e). These interesting observations suggested that HR electrode geometry, with a hole in the middle, encourage the diffusion of the charge carriers in a much better fashion than the traditional disk geometry, by providing a seamless electrode/ electrolyte interface and proper utilization of active electrode surface area, hence the charge injection.

##### In vitro cytotoxicity test

Having excellent electrical and electrochemical properties is not sufficient for implants, as they should also be free of toxic molecules. Therefore, the viability of SH-SY5Y neuroblastoma cells was assessed after 1, 3, and 7 days of culture with the microfabricated HR microelectrode array using Live/Dead images (Figure 5a). Figure 5b illustrates the quantitative analysis of the percentage of living cells, where the microfabricated HR electrode did not affect the viability of SH-SY5Y cells, as they presented a non-significant viability compared to control that was higher than 90% for all days of culture. Their excellent cytocompatibility was also verified using the MTT assay, which is used to measure cellular metabolic activity as an indicator of cell viability, proliferation and cytotoxicity. Figure 5c showed that SH-SY5Y cells cultured with the microfabricated HR microelectrode arrays showed excellent proliferation comparable to control cells and significantly higher in days 3 and 7. These cells viability and proliferation results showed that the microfabricated HR microelectrode arrays did not release any toxic molecules during a week-long culture with SH-SY5Y neuroblastoma cells, and thus should be safe to be employed for *in vitro* and *in vivo* applications.

**Figure 5.**
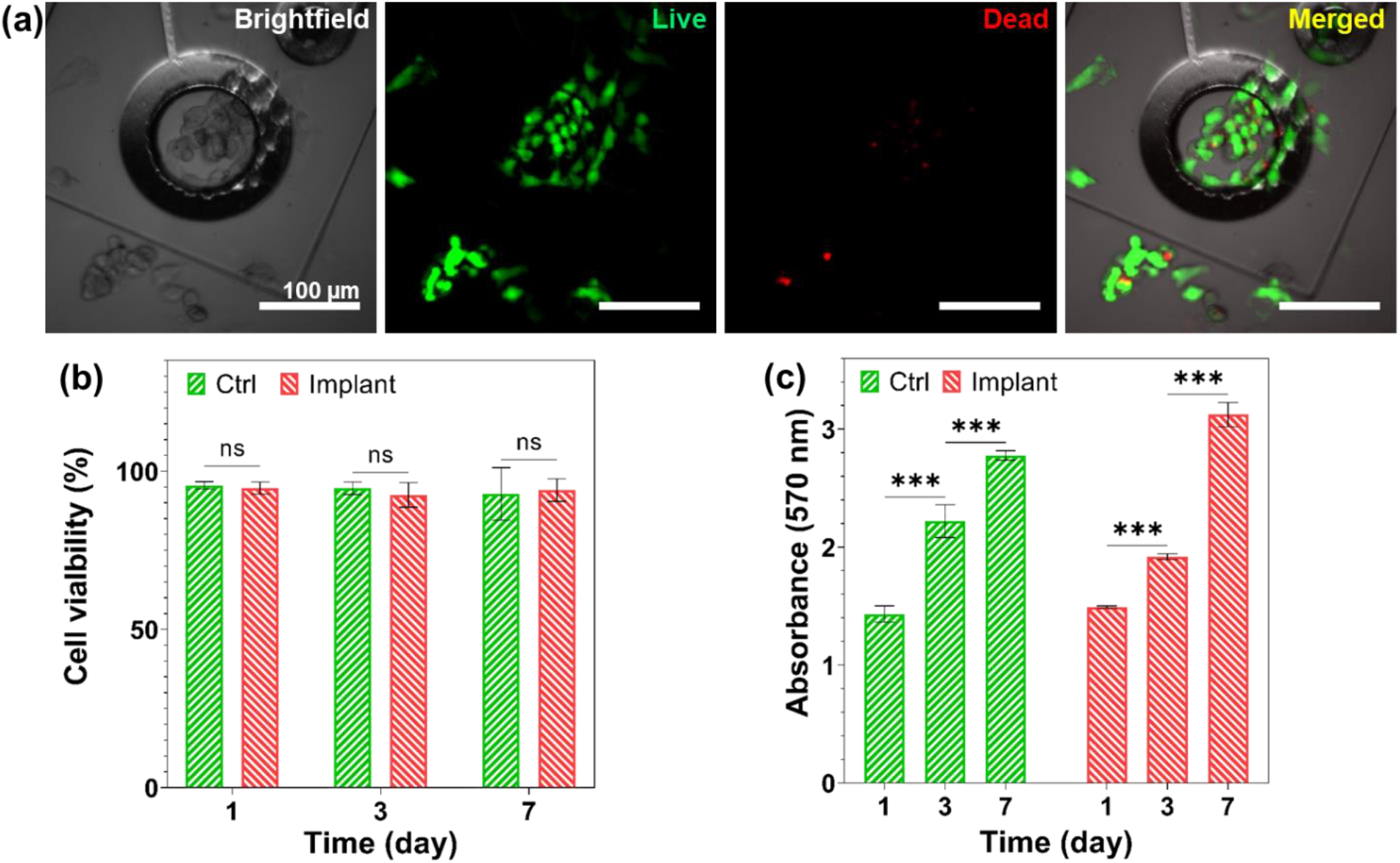
In vitro cytotoxicity assessment of flexible HR microelectrode array. A) Live/dead images of SH-SY5Y cells on day 7. B) Quantified viability of SH-SY5Y cells cultured for 7 days using a Live/Dead Cell Viability Assay (n=6). C) Quantified proliferation of SH-SY5Y cells cultured for 7 days using an MTT Cell Proliferation Assay (n=6). All data are expressed as mean ± standard deviation. Significance is indicated as ***(p < 0.001).

##### Spontaneous electrophysiological activity in 3D neuronal networks

To assess the ability of the HR microelectrodes device to monitor the neural activity of a neuronal system, spontaneous electrical recordings were performed on 3D *in vitro* neuronal cultures at an early age of cell culture DIV 23. Hippocampal neurons, dissociated from E18 Sprague Dawley rat were first entrapped in multi-layered silica beads, and seeded through the HR microelectrodes probe, enabling the growth of neural cells in the 3D matrix and the formation and maturation of a neural network around the HR microelectrode (Fig. 6 a-c).^63-64^ As shown in the Figure 6a & b, we designed the electrode architecture and position, specifically in such a way that it offers multidirectional sensing of neuron activities within the 3D cultures, from both front and back-end of the HR microelectrode. We experimentally evaluated the HR microelectrode performance by recording extracellular action potentials (spikes) and lower frequency burst activities, which is expected for networks from Glutamatergic and GABAergic neurons in 3D cell cultures.^64^ Comparison with control experiments using Disk-type microelectrodes coupled to 3D cell cultures were also carried out.

**Figure 6:**
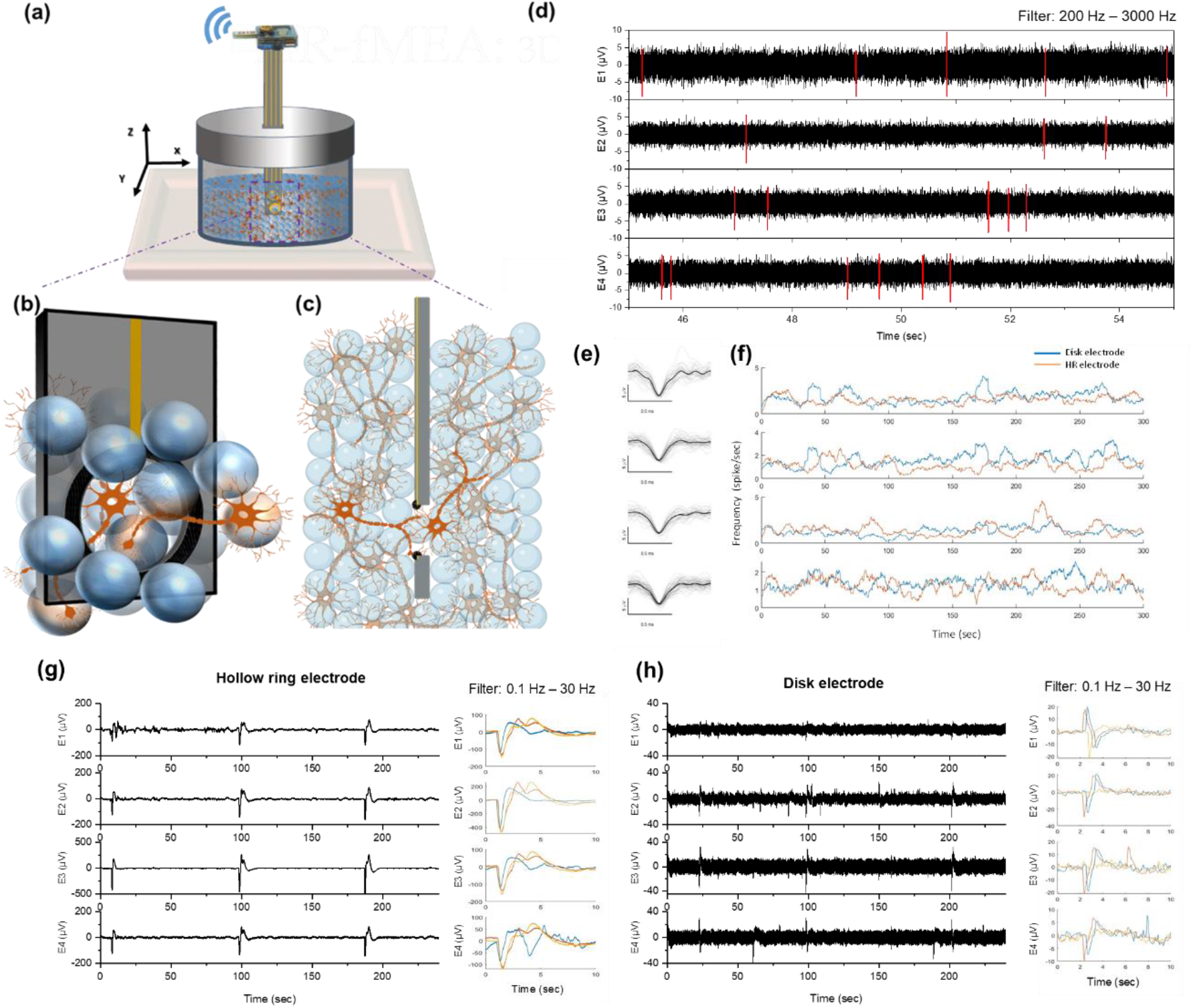
(a) Schematic of *in vitro* neural cell culture monitoring in detail. HR type microelectrode arrays were assembled vertically (z-direction) within the cell culture chamber, composed of silicon beads seeded with rat embryonic (E18) hippocampal neurons and connected to a wireless MCS (W200-HS4) recorder. (b) and (c) are the schematics describing how the HR type electrode interacts with the cells within the cultures, from a closeup and cross-sectional viewpoints respectively. The raw data, obtained from 4 electrodes corresponding to the neural cell recordings were analyzed by separating them into two distinct signals. (d) and (e) Spikes: wireless recording of spontaneous single unit activities, where the raw data was filtered using a bandpass filter 200 Hz – 3000 Hz (d). Red color lines represent the single unit activities and the corresponding (e) panel showing the extracted waveforms. (f) Mean single-unit frequency (spike/sec) of the HR (in red) and the Disk type (in blue) microelectrodes, shown in selected day of culture: DIV 23 for 300 seconds. (g) and (h) Burst activities: monitoring of low frequency events within the 3D neural network responses using filter 0.1 Hz to 30 Hz, and the corresponding right panels showing the burst waveforms of HR and Disk type microelectrodes from each electrode (E1 to E4).

First, the neural activity of neuronal networks was successfully recorded in all reported experiments using both of HR and Disk-type microelectrodes. Examples of spiking and burst activities are presented in Fig. 6 d–h. Burst activity is characterized by the development of quasi-synchronous activity and network dynamics composed mainly of network bursts (i.e., electrical events that involve most of the recording channels, which are displayed by low-pass filtering (0.1–30 Hz) of the voltage trace (Fig. 6 g-h). Bandpass filtering (200–3000 Hz) of the original trace reveals the action potentials of single neural cells (spikes) (Fig. 6 d-e).

An overlay of spontaneous action potential waveforms captured by both HR and Disk microelectrodes is shown in Fig. d-e and Fig. S5 respectively. HR microelectrodes presented a similar level of spiking rate (1.6 spike/sec) than disk-type electrodes (1.7 spike/sec), while spike amplitudes and durations were not significantly different between the two. By looking closer, the HR electrode exhibited lesser noise in general and from the features of action potentials derived from single neuron activities, we noticed mainly negative spike waveforms having peak amplitudes up to 8 μV. As shown in the literature, they represent the action potentials propagating along axonal branches that are often neglected while measuring with bigger size electrodes.^65^ This result suggests that the recording strength on HR electrodes is higher than Disk-type electrodes with lowering of noise leading to better signal to noise ratio (SNR). Indeed, lower electrical impedance in HR microelectrode (Fig.3) improves the SNR through a combined effect of lower noise levels, predominantly thermal noise.^66^

We also observed in our recordings that there are quasi-synchronized events that typically characterise network dynamics composed mainly of network bursts (Fig. 6 g-h). The presence of bursts indicates the existence of neural network connectivity and they typically last a few hundreds of milliseconds to seconds, which is expected for functional and mature networks. For disk-type recordings, these bursts are an amplitude of about 20-30 μV and a duration of about 500-1000 ms (Fig. 6h). Besides, Figure 6g shows an example of bursts activities (same batch of cell culture) recorded with the HR microelectrodes from a 3D network at the same age of its development (i.e., =DIV 23). From a qualitative observation, it can be noticed that the signature of the network dynamics (i.e., bursts) show a distinct mode of firing, where the bursts are characterized by a higher amplitude (i.e., about 100 to 500 μV),) and a longer duration (up to 5 s).

Typically, it is assumed that large electrodes (SA > 2000 μm^2^) are well suited for recording population-wide filed potentials, while small electrodes (SA < 500 μm^2^) are more suitable for detecting multi-unit spiking activities or individual neural cells (single-unit activity).^65^ Unlike traditional Disk type microelectrodes, our HR architectures having the surface area of 450 μm^2^ - 1400 μm^2^ with a hole of 40 – 100 μm in the middle, demonstrated that they can capture both sub-cellular features of neuronal signals (spikes) and propagation of high-resolution burst signals at lower frequencies.

## 4. CONCLUSIONS

In this study, a novel Hollow Ring type neural interface electrode has been designed and fabricated. The flexible ring electrode consisted of two layers of Parylene C with gold electrodes sandwiched in a ring-shaped geometry. Thanks to its unique architecture, ring-shaped electrodes enable easy and reliable access of the electrode to 3D neural networks with minimal pressure on the biological tissue, while still providing improved electrical contact with cells. The ring electrodes have a couple advantages from the design-performance relationship point of view. Among the planar microelectrode geometries studied, the ring design appears to be a superior geometry for performing electrochemical measurements. The interfacial impedance of the electrode is dramatically reduced compared to metal and disk-type electrodes. In addition, proposed electrode ring geometries, have well-like structures for millisecond biphasic stimulating microelectrodes. The results from voltage excursion experiments showed that electrode geometries with a ring hole yield a significant increase in charge injection limit compared to the traditional planar structure of similar dimensions. Electrophysiological investigations further confirmed the hypothesis that ring type interface electrode indeed provide reliable and improved neuroelectrical interface. The results of the neural recordings from three-dimensional neural networks demonstrated that differences in low frequency bursts could be distinguished using the Ring-type electrode configuration. Also, we showed that the SNR of the single-unit activities recorded by the Hollow Ring electrode was higher than those from Disk electrodes. In summary, our results suggest the great potential of the HR electrode design for developing next-generation microelectrodes for applications in neural interfaces used in physiological studies and neuromodulation applications.

## ACKNOWLEDGMENT

The authors acknowledge fundings from the Agence Nationale de la Recherche (ANR-19-CE19-0002-01). This work was supported by French RENATECH network.

## Supplementary Information

The following files are available free of charge: Experimental details, Table S1 and Fig. S1−S5

## Supporting information

**Figure S1:**
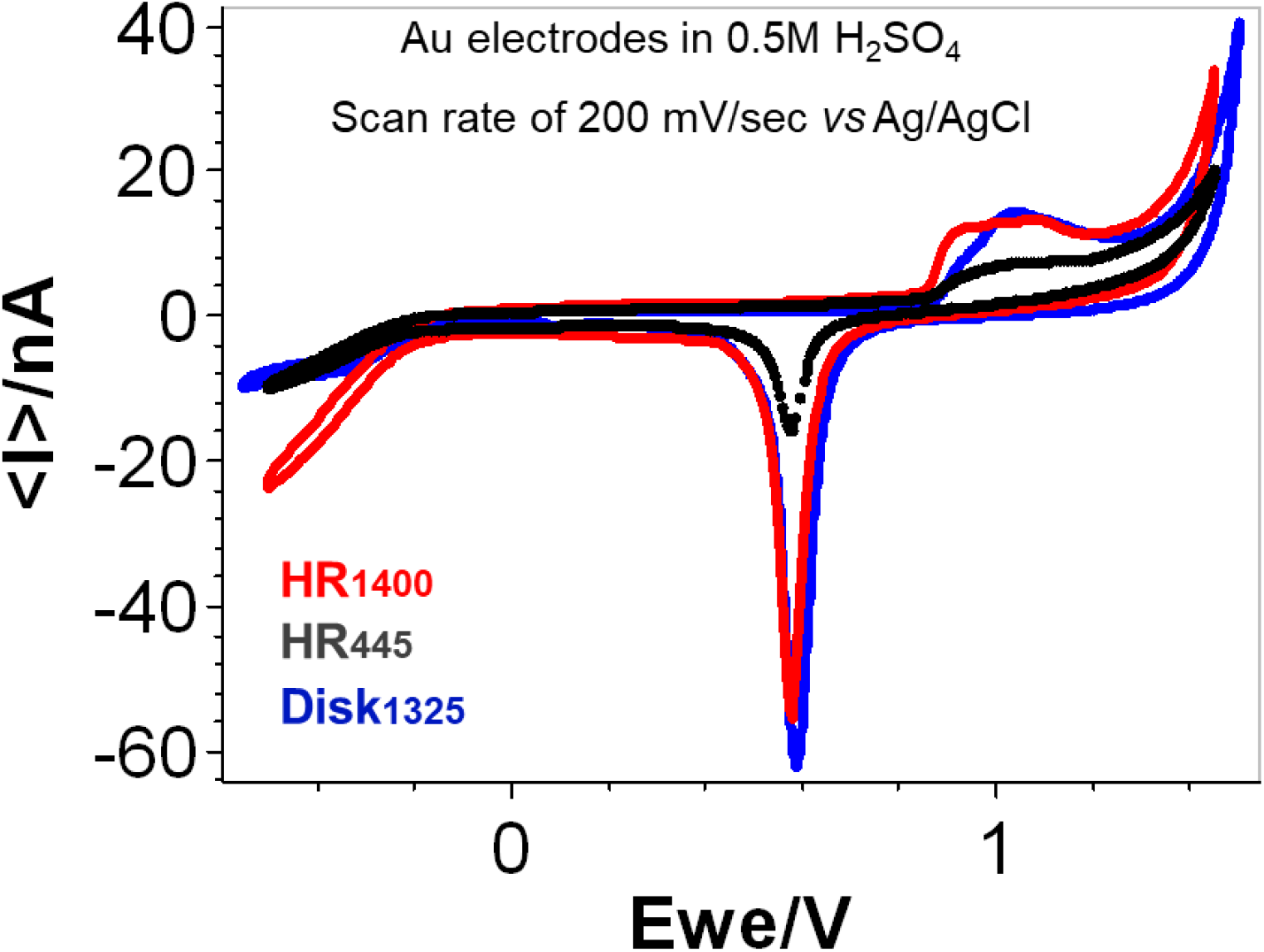
(a) Representative cyclic voltammetry graphs of Hollow Ring (HR_1400_ and HR_445_) and Disk type gold electrodes in 0.5M H_2_SO_4_ solution at a scan rate of 200 mV/sec *vs* Ag/AgCl reference.

**Figure S2:**
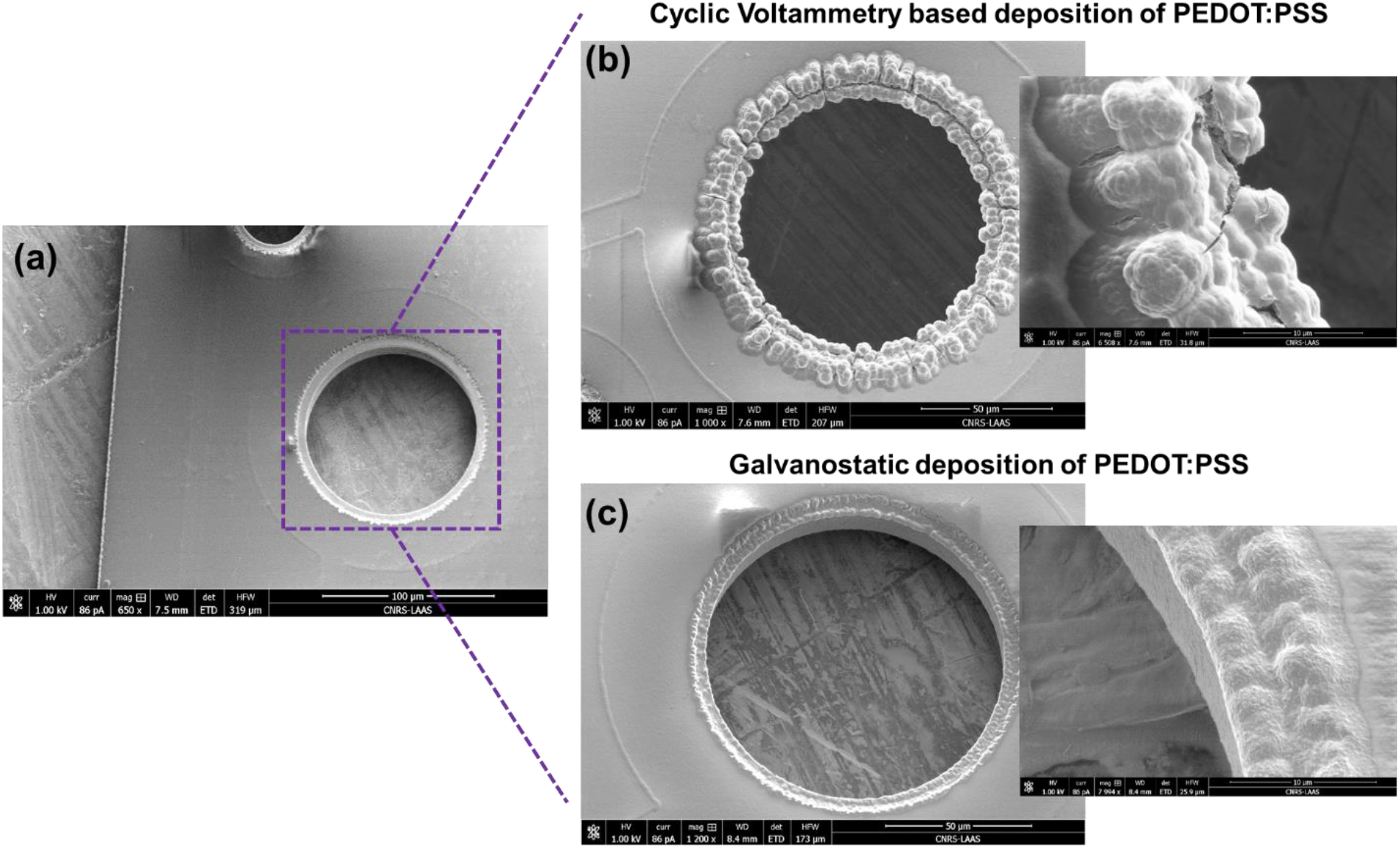
SEM micrographs of HR-fMEA, (a) before and (b & c) after the PEDOT:PSS deposition. For CV based deposition, every electrode was subjected to one CV scan cycle between -0.7 V and 0.9 V, at a scan rate of 10 mV/s vs Ag/AgCl reference electrode. Whereas, in the case of galvanostatic deposition technique, the PEDOT: PSS was deposited by applying a small current of 12.5 nA, with a constant charge density of 4 nC/μm^2^.

**Figure S3:**
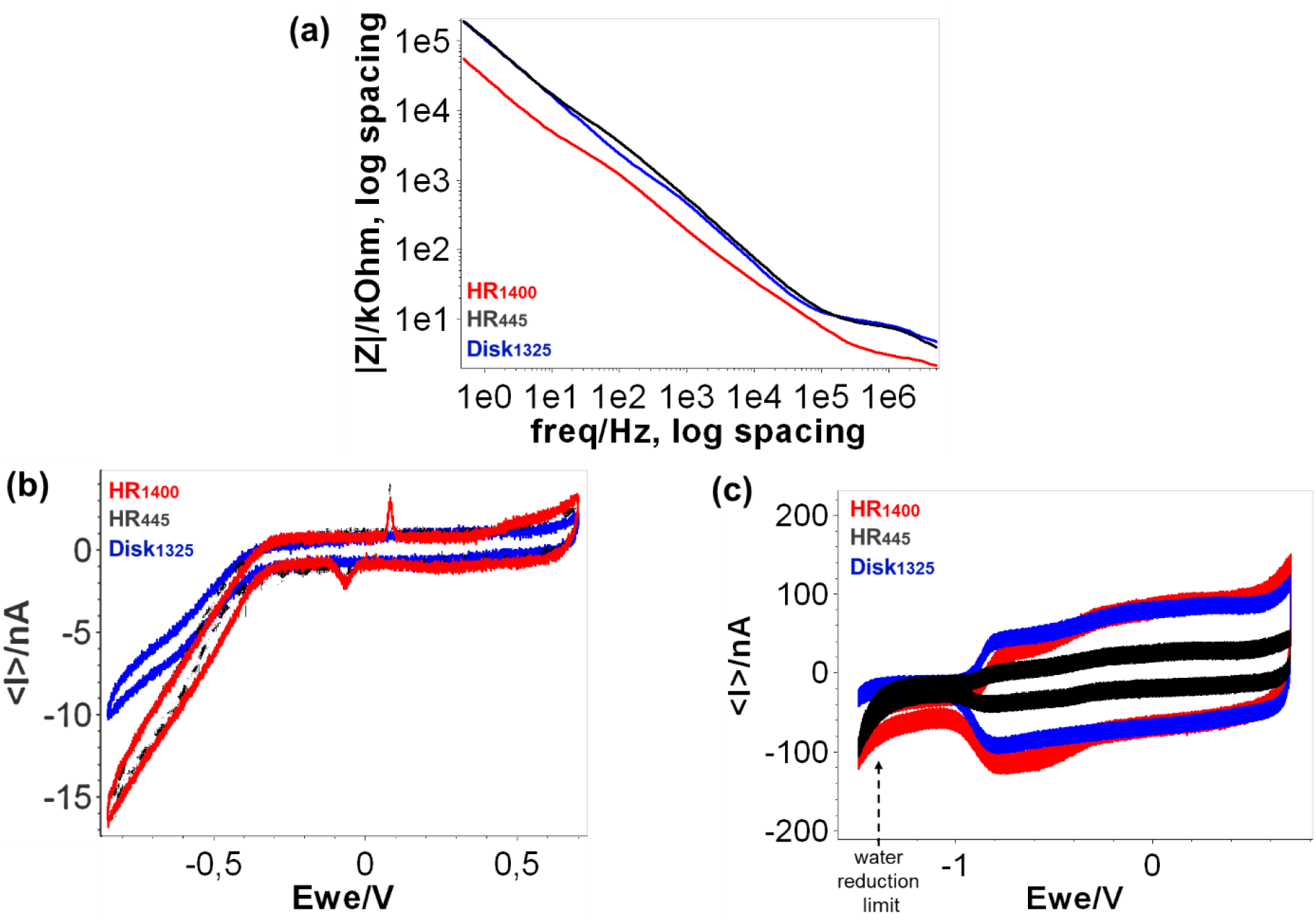
(a) Bode plots of HR and Disk type gold electrodes representing the |Z| vs frequency over a frequency range of 0,1 Hz to 5 MHZ in PBS at 0V vs Ag/AgCl reference electrode. Comparison of electrochemical windows (b) before and (c) after PEDOT:PSS deposition, and the respective water reduction potential by cyclic voltammetry. PBS buffer was used at a scan rate of 200 mV/sec *vs* Ag/AgCl reference.

**Figure S4:**
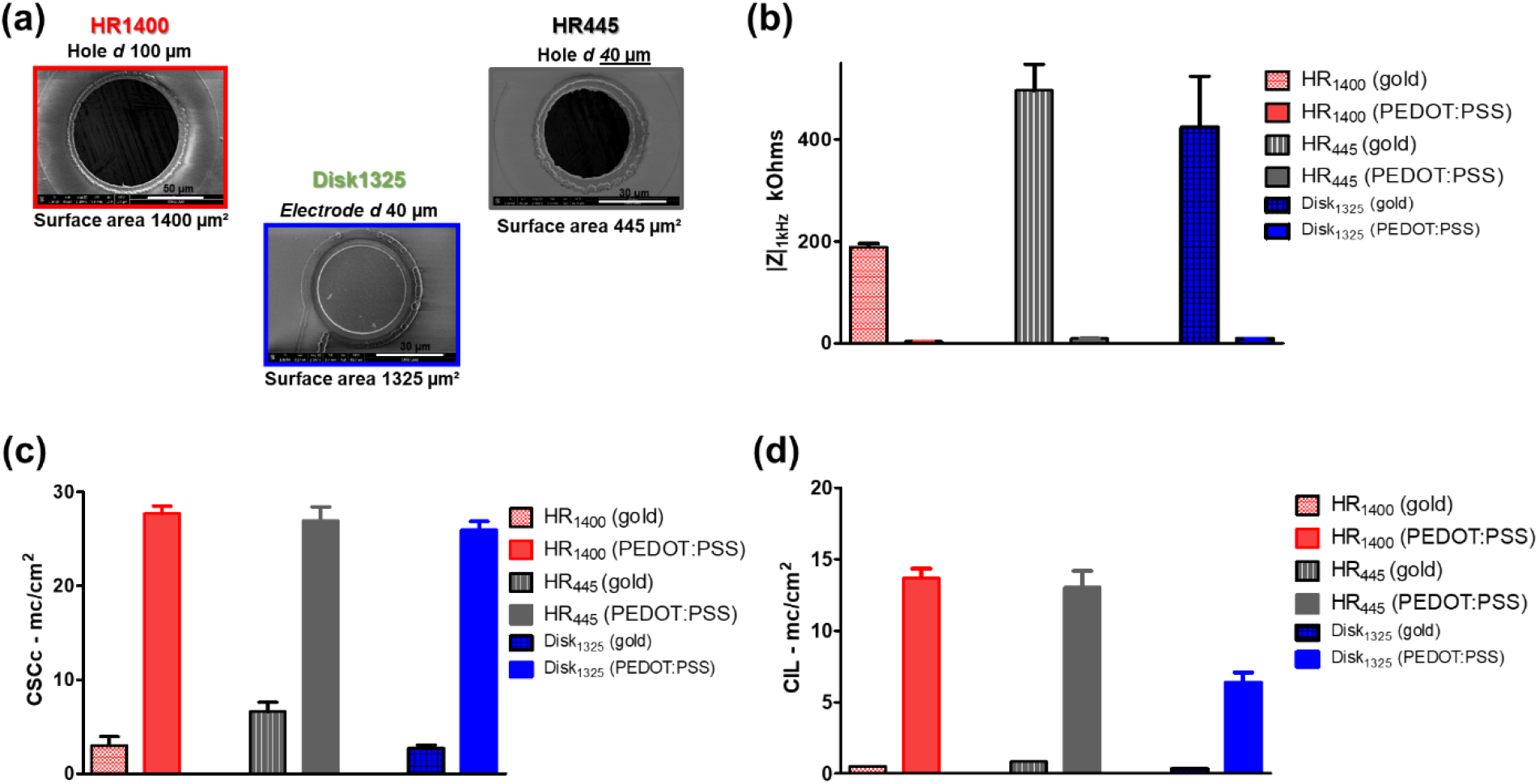
(a) SEM images of Hollow Ring type and Disk type electrodes. The average (b) EIS at 1KHZ, (C) CSCc and (d) CIL values of HR and Disk type electrodes, before and after PEDOT:PSS coating.

**Figure S5:**
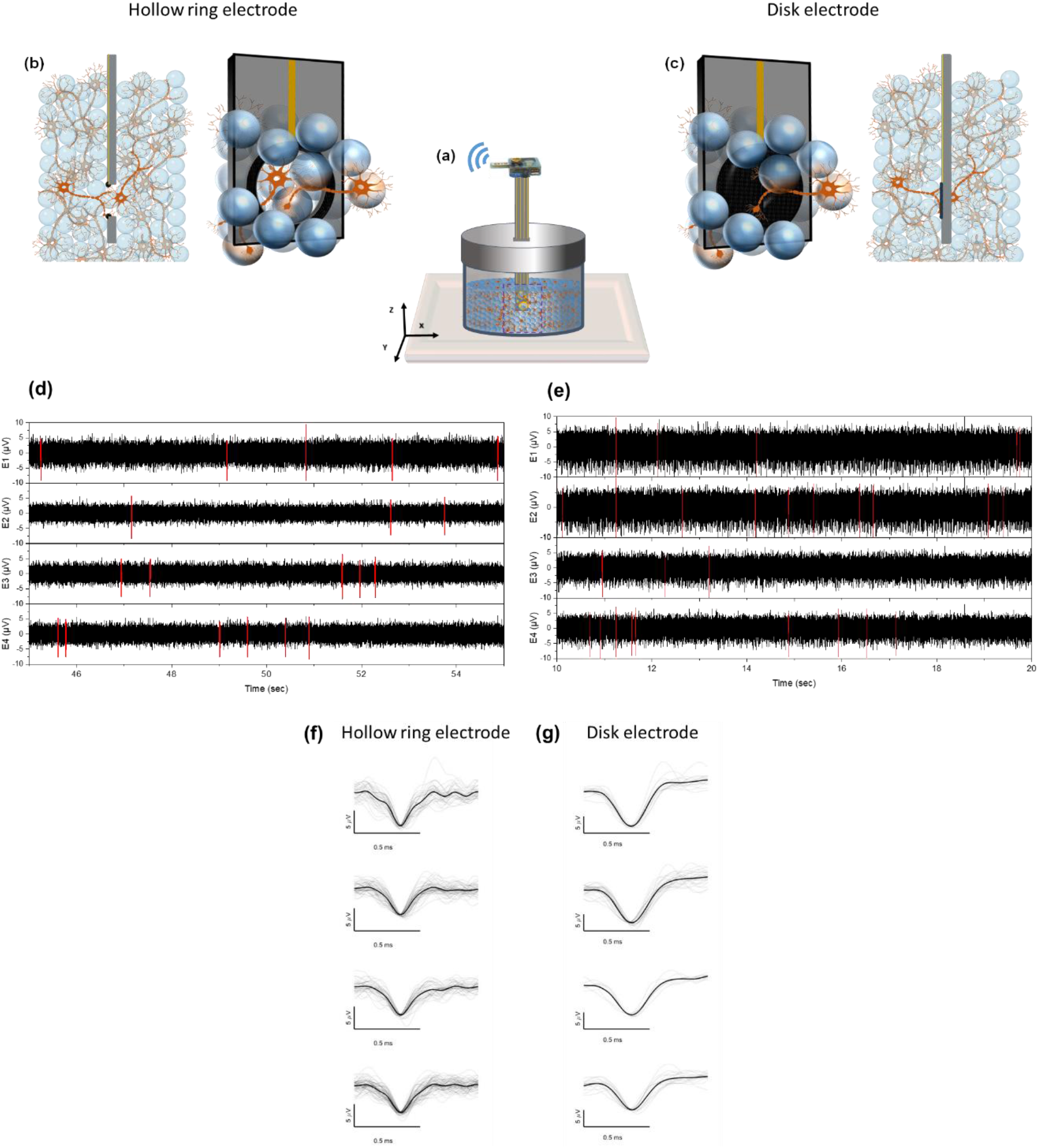
(a) Schematic of *in vitro* neural cell culture monitoring in detail. HR and/or Disk type microelectrode arrays were assembled vertically (z-direction) within the cell culture chamber, composed of silicon beads seeded with rat embryonic (E18) hippocampal neurons and connected to a wireless MCS (W200-HS4) recorder. (b) and (c) are the schematics describing how the HR and Disk type electrode interacts with the cells within the cultures, from a closeup and cross-sectional viewpoints respectively. The raw data, obtained from 4 electrodes corresponding to the neural cell recordings were analyzed by separating them into two distinct signals. (d) and (e) Spikes: wireless recording of spontaneous multiple unit activities, where the raw data was filtered using a bandpass filter 200 Hz – 3000 Hz. Red color lines represent the single unit activities and the corresponding (f) and (g) panels showing the single-unit waveforms from four electrodes (E1 to E4).

